# Free Aerial Imagery as a Resource to Monitor Compliance with the Endangered Species Act

**DOI:** 10.1101/204750

**Authors:** Jacob Malcom, Tiffany Kim, Ya-Wei Li

## Abstract

Compliance monitoring is an integral part of law and policy implementation. A lack of compliance monitoring for the Endangered Species Act (ESA), driven in part by resource limitations, may be undercutting efforts to recover threatened and endangered species. Here we evaluate the utility of freely available satellite and aerial imagery as a cost-efficient component of ESA compliance monitoring. Using data on actions authorized by the U.S. Fish and Wildlife Service (FWS) under section 7 of the ESA, we show that approximately 40% of actions can be found in remotely sensed imagery. Some types of actions, such as residential and commercial development, roadwork, and forestry, show substantially higher observability. Based on our results and the requirements of compliance monitoring, we recommend FWS standardize data collection requirements for consultations; record and publish terms and conditions of consultations; and encourage their staff to use technology such as remotely sensed data as a central part of their workflow for implementing the ESA.

## Introduction

The effectiveness of laws and policies is a function of their strength and the degree of compliance: a superb law with low compliance may achieve less than a mediocre law with high compliance. Policymakers cannot accurately assess the effectiveness of a given law or policy without understanding the extent and consequences of noncompliance by both regulating agencies and regulated parties (Farber, 1999; Keane et al., 2008). Thus, monitoring and enforcement are integral to policy creation, implementation, and learning.

The U.S. Endangered Species Act (ESA) is considered the premiere legislation for protecting threatened and endangered wildlife. Researchers estimate the law has prevented the extinction of over two hundred species (Scott, Goble, Svancara, & Pidgorna, 2006). Even with its successes, current implementation of the ESA may be insufficient to protect biodiversity in the face of increasing threats. Less than 2% of all listed species have been delisted due to recovery, and from 1990-2010, half of listed species were still declining, 35% were stable, and only 8% were improving (Evans et al., 2016). The causes of relatively low recovery rates are unclear. One likely explanation is funding: expenditures on listed species tend to be far below what scientists predict is needed for recovery (Gerber, 2016). Another possibility is that compliance with the ESA may be low: regulated entities may continue to harm species and their habitats faster than recovery can occur, but this hypothesis is untested.

Section 7 of the ESA (Box 1) is arguably the law’s strongest regulatory provision for many listed species, at least on paper (Malcom & Li, 2015). It requires federal agencies to use their authorities to conserve threatened and endangered species, including by consulting with the U.S. Fish and Wildlife Service (FWS) and/or the National Marine Fisheries Service (NMFS) on federal actions that may harm those species. Over 10,000 projects are evaluated by FWS in section 7 consultations each year (Malcom and Li, 2015), which means >10,000 opportunities for permittees to avoid or minimize their impact on ESA-listed species and >10,000 opportunities for FWS to encourage or require conservation actions that can help advance recovery. But it also means >10,000 chances for permittees to fall short of their duties, i.e., be out of compliance and potentially harming species more than allowed.

#### Box 1: ESA Section 7 consultation overview

Section 7 requires all federal agencies to cooperate to conserve endangered species. Section 7(a)(1) directs federal agencies to use their authority to conserve threatened and endangered species. Section 7(a)(2) requires federal agencies to avoid jeopardizing the existence of threatened or endangered species or the destruction or adverse modification of their designated critical habitat. To comply with Section 7(a)(2), federal agencies must consult with the U.S. Fish & Wildlife Service (FWS) when they intend to carry out, permit, or fund an action that might harm ESA-listed terrestrial or freshwater species. The applicant generally begins with an “informal consultation,” but if an action is “likely to adversely affect species or critical habitat,” a formal consultation is required (Handbook 1998). In formal consultations, FWS must issue its biological opinion that determines whether the action may jeopardize the species or destroy or adversely modify critical habitat. If neither of these are likely to occur, the action agency may proceed with its project under an incidental take statement (ITS) prepared by FWS that details the amount of “take” the agency is allowed, often expressed as a number of individuals or area of habitat. Any action exceeding this level of take would trigger reinitiation of the consultation process (and violate section 9's prohibition on unauthorized take). The ITS also includes mandatory “reasonable and prudent measures” to minimize the level of take, and stipulates monitoring requirements to, ostensibly, ensure that the action follows the description in the consultation.

Given the volume of actions authorized under section 7, compliance monitoring is a critical aspect of implementing the ESA. We need to know if an applicant cleared only the 5 acres of Florida panther (*Puma concolor coryi*) habitat they proposed to clear, or if they actually cleared ten acres and hoped nobody would notice. Did the applicant install silt fences along the highway they were widening near a stream with blackside dace (*Phoxinus cumberlandensis*)? Was ecological restoration in one area sufficient for offsetting habitat destruction elsewhere? Formal consultations require submitting regular monitoring reports, but whether FWS personnel have time to review those reports or independently verify their content is questionable. For example, a 2009 Government Accounting Office (GAO) study found that 63% of assessed consultation files did not have all the required reports available, including almost 40% that had no reports at all. Most FWS managers rely on individual biologists to carry out informal monitoring (e.g., telephone calls, emails), which hinders transparency and causes problems when institutional knowledge is lost as a result of personnel turnover (GAO 2009). In a worst-case scenario, the GAO describes a consultation for which one agency did not submit any of the required monitoring reports until fifteen years after the action date. At that point, the population of the listed species declined from 1400 individuals to “none found at all” (GAO 2009). Furthermore, the lack of monitoring results in inadequate information about the effectiveness of conservation measures stipulated by FWS to minimize take (Totoiu, 2011). These shortcomings mean that FWS could authorize too much incidental take or specify ineffective conservation measures, even when doing so would jeopardize a species’ existence.

Remotely sensed data is one possible resource that can aid environmental compliance monitoring (Purdy, 2006; de Leeuw et al., 2010). These data have been used in many studies of tropical deforestation (e.g., Skole and Tucker, 1993; DeFries et al., 2002; Baccini et al. 2012), but there have been few attempts to use them for direct enforcement of specific environmental laws. One exception is found in Australia, where multiple state agencies have been using satellite imagery to effectively monitor site compliance and hinder illegal deforestation for over a decade (Purdy, 2010). In other cases, personnel from an NGO used satellite and aerial imagery to show that the State of Texas had been dramatically under-reporting habitat destruction from oil field development (Li, Shepard, & Male, 2013), and nighttime satellite imagery was used to evaluate compliance with sea turtle light ordinances in Florida (Anderson, Nuernberger, Yamamoto, & Sutton, 2013). But the rarity of these cases suggests there may be ample room to use remotely sensed data to improve monitoring and enforcement. A critical outstanding question is how often remotely sensed data can be used to monitor compliance with section 7 of the ESA.

Here we report on the utility of satellite and aerial imagery as a key resource for ESA section 7 compliance monitoring. We use geographic coordinates for hundreds of consulted-on actions, from a database of all section 7 consultations from 2008-2015, to show that many of these actions can readily be identified in freely available satellite and aerial imagery. If this data source and approach are combined with regulatory data that specify FWS’s expectations of or limits on these actions (e.g., area to be disturbed), any interested party—government personnel, NGOs, or private parties—can contribute to ESA compliance monitoring.

## Methods

As a first step to evaluating the utility of satellite and aerial imagery as a component of section 7 compliance monitoring, we queried a database of consultations described by Malcom and Li (2015). The database is a curated version of FWS’s Tracking and Integrated Logging System (TAILS) database, and contained 88,290 section 7 consultations recorded by FWS biologists from January 2008 to April 2015 at the time of our query. Rather than selecting a random sample from among all consultations, we first filtered out consultations for which FWS personnel did not record coordinates for the action; we have very little chance of finding those actions. Next, we narrowed the pool of possible consultations to those for the 20 most common action types for informal and formal consultations using the Section 7 Explorer web application (Malcom and Li 2015; https://cci-dev.org/shiny/open/section7_explorer/). We reasoned that it will be more useful to have more samples from common action types than just one or two samples of rare action types. From this reduced pool we collected a random sample of informal and formal consultations. If the sample for an action type included fewer than three consultations, we randomly selected additional consultations from within that category to ensure a minimum sample size of three per category.

To determine whether we could see the effects of each action in remotely sensed images, we navigated to the action coordinates in Google Earth Pro (GEP). Using the “Historical Imagery” view, we identified the image nearest in time to the consultation end date (after which the project could proceed) to establish a baseline. We then stepped through the sequence of subsequent available images and watched for any signs of relevant activity at or around the coordinates. For example, if the action type was “development” then we watched for any signs of residential or commercial development. We recorded the date of the first image showing relevant activity (if any) and the date of the first image in which that activity appeared to have been completed or ceased. If no activity was visible or the end of the activity was unclear, we examined every image available up to May 2016. We found it was sometimes necessary to look up to 2.5 km from the given coordinates to find the likely action, e.g., when a land management agency reported the coordinates at the center of their district rather than the actual action site. To get a sense of the types of conversions we found, we recorded the observed change in habitat type (e.g., from a “natural” habitat type to developed land).

We also sought to test our assumptions about the observability of different types of actions. To do so, two authors—JWM and TLK—independently scored all 448 types of actions recorded in TAILS according to whether they thought such actions would be observable (likely to be observable = 1, unknown = 0.5, unlikely = 0). We identified discrepancies between our scores, discussed the reasoning for the discrepancies, and settled on a consensus observability for each action type. We qualitatively compare our expectations to what we observed in the imagery.

All data underlying this paper, code for the analyses, supplemental information, and additional content (e.g., example images) are archived at the Open Science Framework (https://osf.io/ane85/; doi: 10.17605/OSF.IO/ANE85). All analyses were done in R 3.3 (R Core Development Team, 2016). The analyses required only basic calculations such as means and proportions. We packaged the full analysis as an R vignette, available from GitHub at https://github.com/Defenders-ESC/compliance_exploration.

## Results

Our dataset included 364 consultations (182 formal and 182 informal) selected from among 42,563 (48.2%) consultations recorded with geographical coordinates by FWS from 2008-2015. This sample constitutes ~2% of formal and ~0.2% informal consultations. The locations of the consultations in our sample generally reflected the distribution of consultations observed by Malcom and Li (2015): formal consultations were more likely to come from the western US, informal consultations more likely from the East (Figure 1). The bias of which FWS offices do or do not record coordinates is apparent for some states, like Washington (*n* ~ 500 formal consultations, many with coordinates recorded) versus Colorado (*n* ~ 500 formal consultations, but few with coordinates recorded). The top consulting agencies in our sample also tended to reflect the top consulting agencies across all agencies (Table 1; for comparison, see Malcom and Li, 2015). The number of actions by type of action in our sample varied substantially between formal and informal consultations (Figure 2). The oldest available aerial imagery for each of the sites evaluated ranged from 1939 to 2005, with an average date of August, 1992 (SI Figure 1).

**Table 1.**
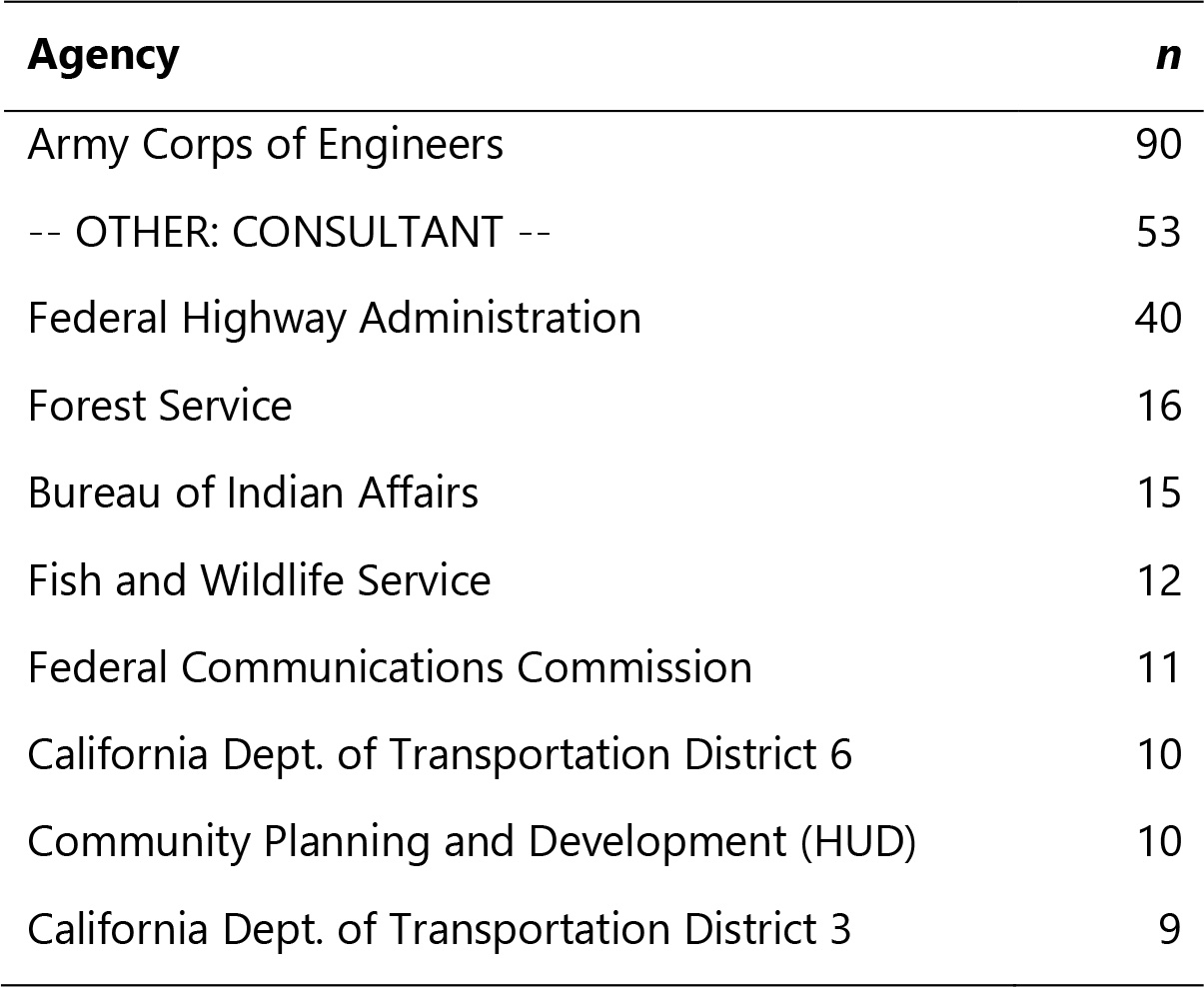
The most common consulting agencies in our sample of consultations.

**Figure 1.**
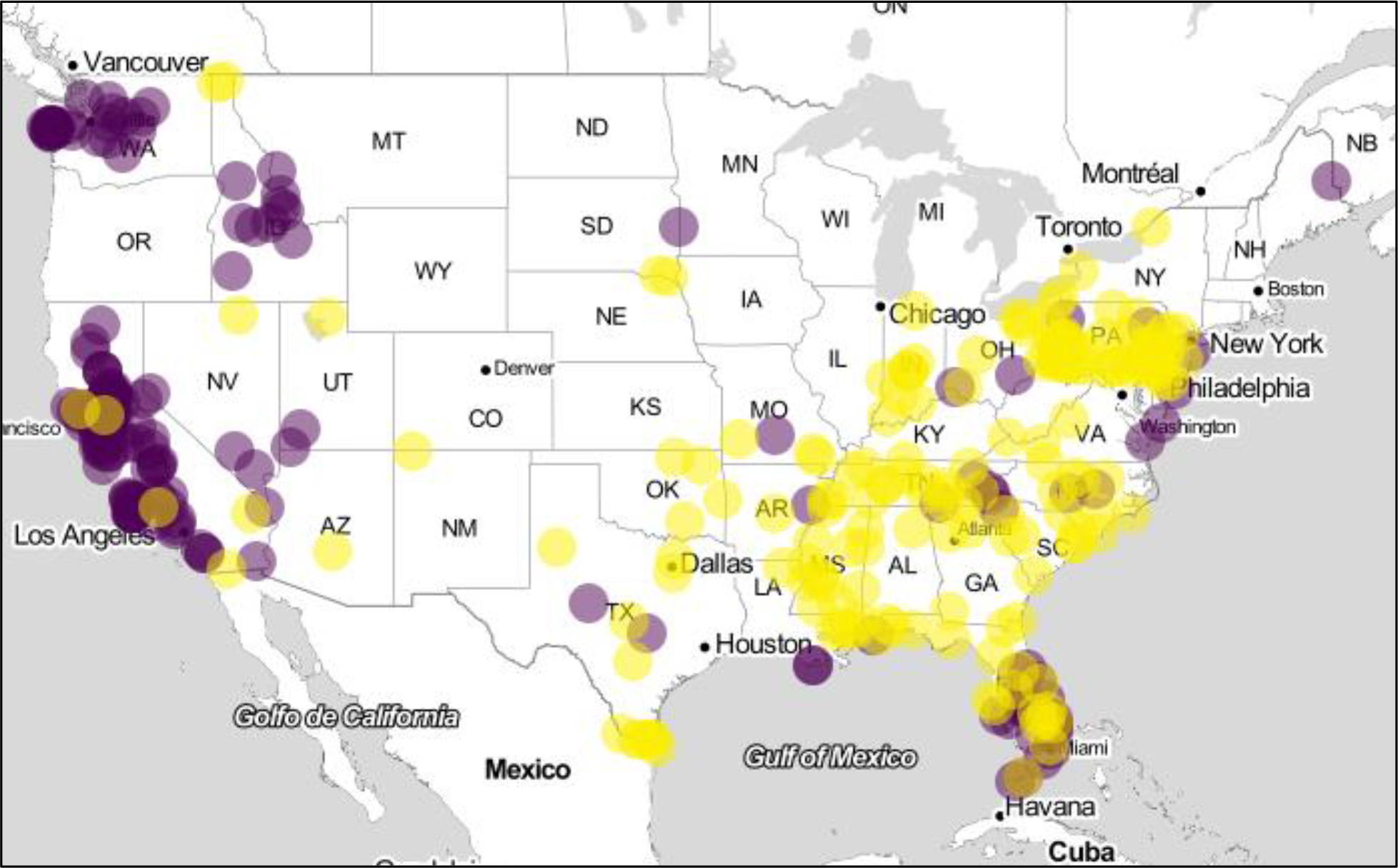
The geographic distribution of consultations in our sample generally reflects the distribution of all consultations. Informal consultations (yellow circles) are more common in the eastern United States and formal consultations (purple circles) are more common in the western United States. Some Fish and Wildlife Service offices tend to not record the geographic coordinates of consultations (e.g., Colorado offices), while others tend to be much better (e.g., Washington state offices). See Malcom and Li (2015) for maps of all formal and informal consultations.

**Figure 2.**
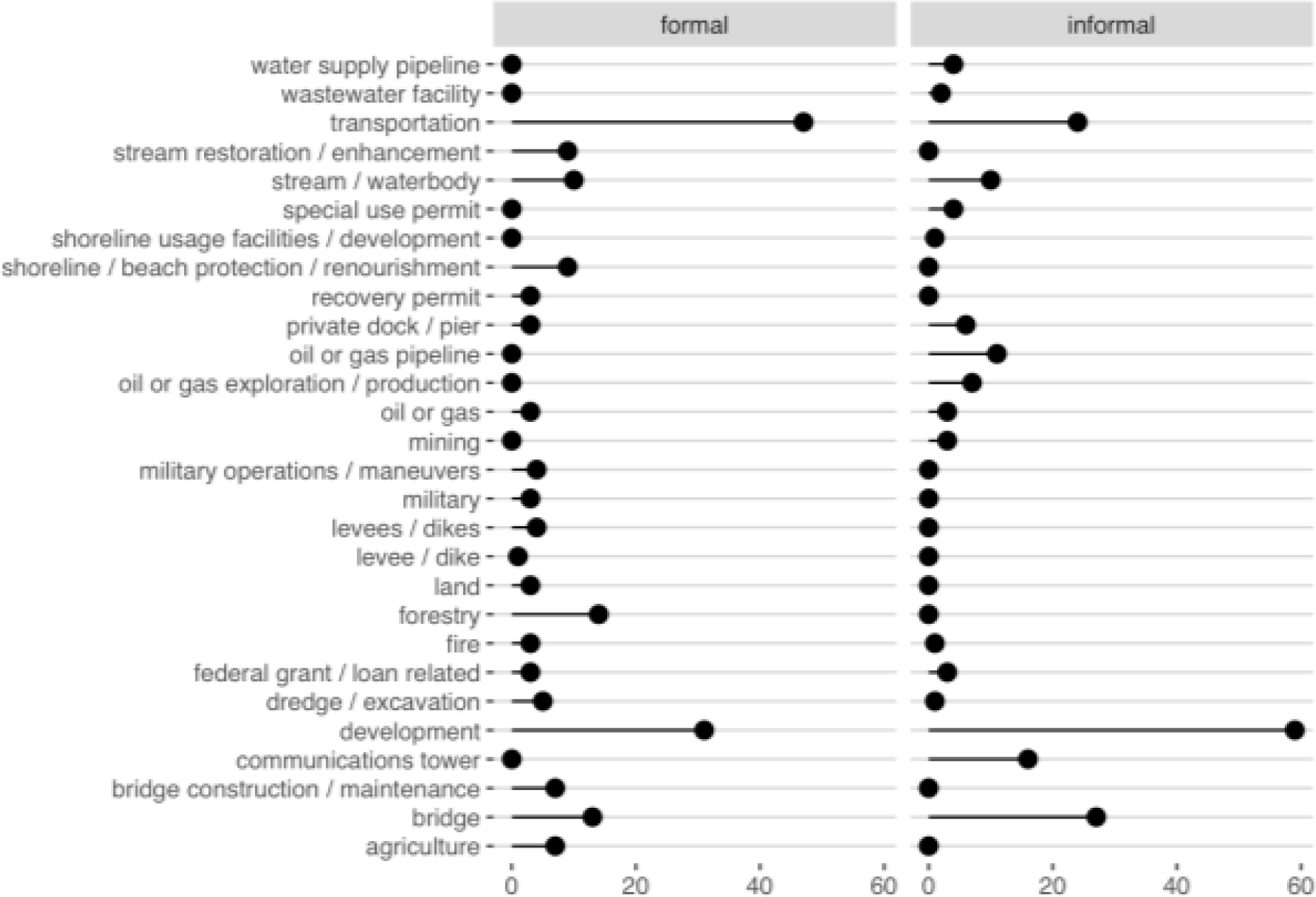
The sample size typically varied between formal and informal consultations for each of the action types we examined. The “lollipops” show the sample size for each action type in the formal and informal consultation types. Because of differences in impacts on listed species, some action types are common in one consultation type but very rare in the other (e.g., communications towers).

Based on the types of actions recorded, we expected formal consultation actions would have observabilities just over 0.5 and informal consultation actions just over 0.65 (SI Table 1). We estimated the observabilities for actions from both formal and informal consultations to be 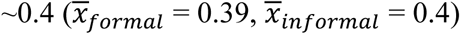. Our overestimate of expected mean observability was caused by our “may be observed” action types more commonly converting to the “not found” category once we looked (Figure 3). Observabilities for different work types varied between zero—we could find no evidence of the action—and one (Figure 4, SI Table 2). The types of observed habitat changes were typically very apparent, with only a small number of samples changing from one “natural” habitat to another (Table 2). A development project in Florida (consultation 41420-2008-F-0112-R001) is an example of natural to development conversion (Figure 5).

**Table 2.** Frequencies of habitat changes found at action sites by consultation type (formal versus informal).

| **Observed conversion** | **Formal** | **Informal** |
| --- | --- | --- |
| agriculture → development | 5 | 13 |
| development → development | 24 | 26 |
| natural → development | 37 | 37 |
| natural → natural | 9 | 0 |

**Figure 3.**
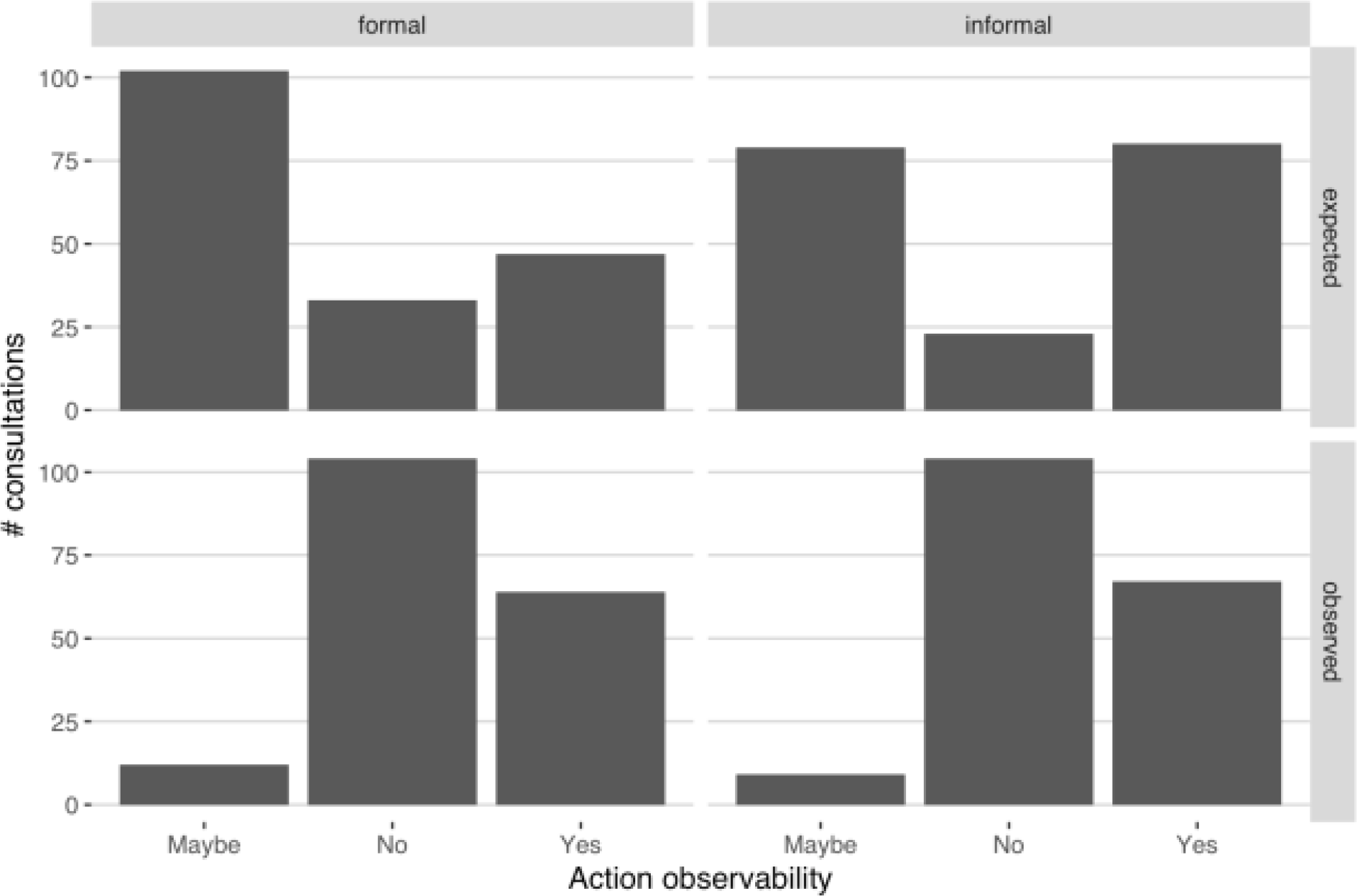
Based on types of actions consulted on, we expected a good bit of uncertainty in observability (top row), but found that uncertain action types tended to not be observable (bottom row). The bars represent the number of consultations we expected to be observable (score = 1), not be observable (score = 0), or perhaps be context-specific (maybe observable; score = 0.5). The top row shows the distribution of expected observabilities given our scores for action types and what our selected consultations covered. The bottom row shows the distribution of observabilities after we had reviewed all 364 consultations in our sample.

**Figure 4.**
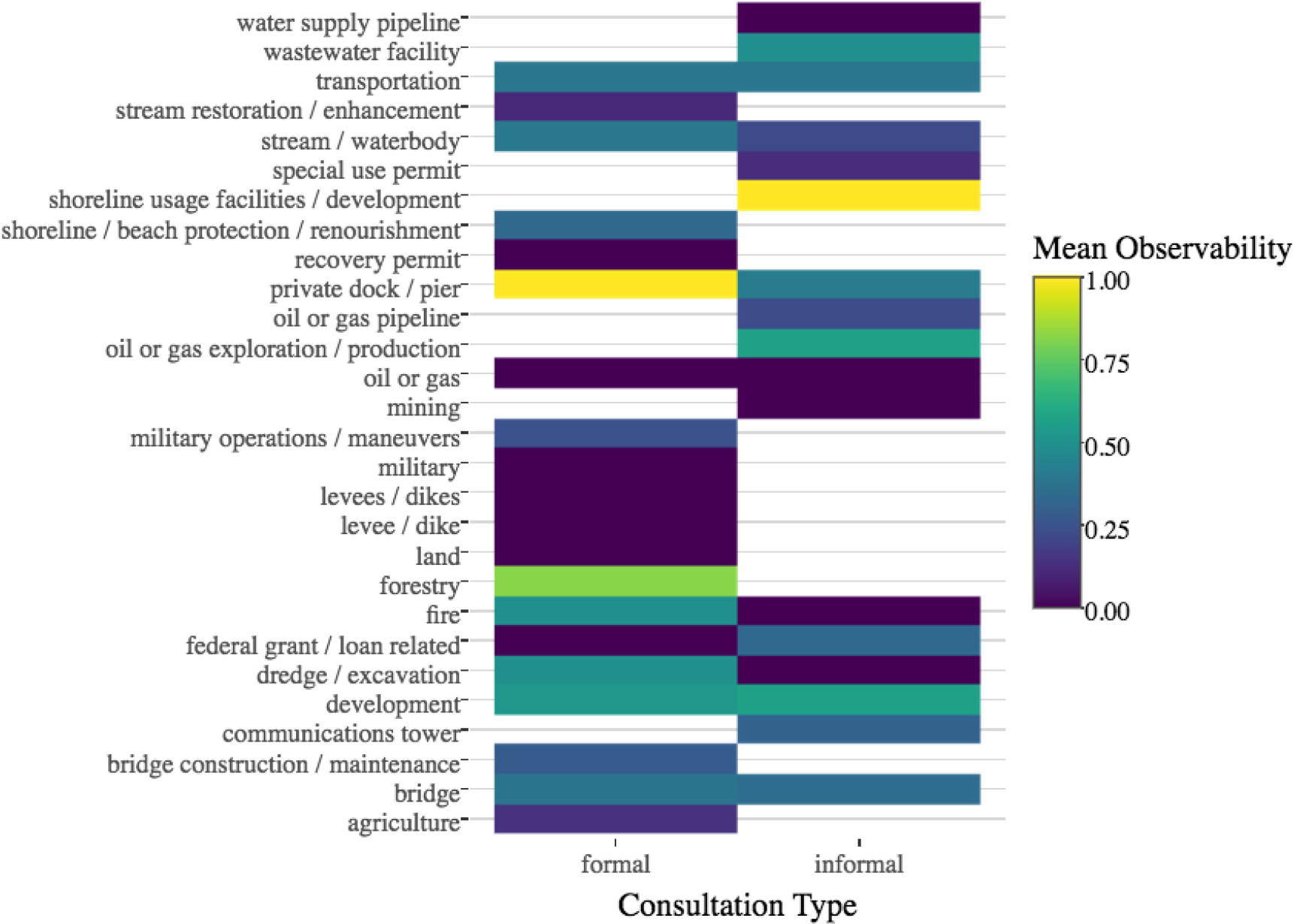
The observability of the 28 action types we surveyed ranged from 0 to 1. A heat map of observabilities by consultation type (formal vs. informal) and action type. Blank entries occur for action-consultation type combinations not in our sample, even if that combination may occur among all consultations.

**Figure 5.**
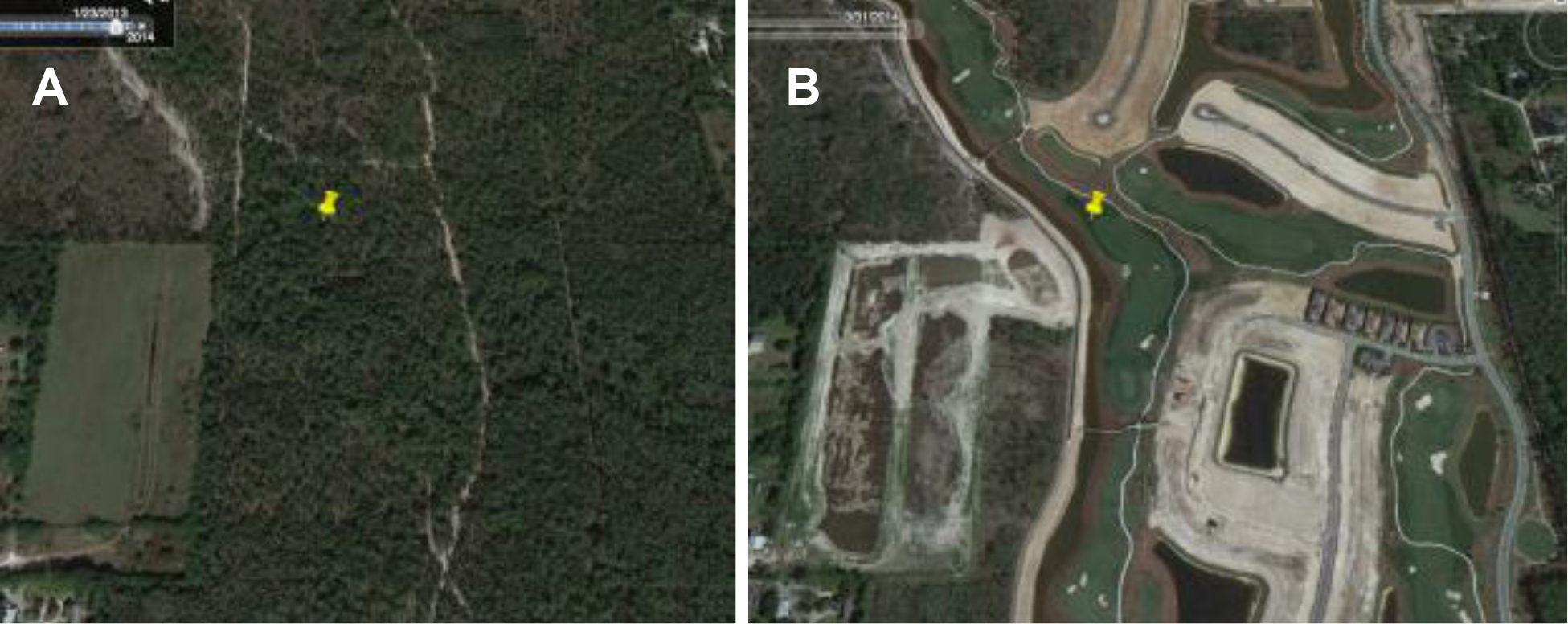
Habitat loss between 2013 (A) and 2014 (B) from a development action that went through formal consultation with the Fish and Wildlife Service (FWS). This site is near Naples, Florida. The consulting agency was the Army Corps of Engineers for a Clean Water Act section 404 permit to dredge or fill wetlands. FWS determined the Florida panther (*Puma concolor coryi*) would be harmed, but its existence would not be jeopardized, as a result of the development.

Using the average observability across work types and assuming FWS had recorded geographic coordinates for all consultations, we expect users might be able to find ~35,300 of the 88,290 consultations in TAILS. (Because 48% of consultations in TAILS do not include coordinates, only ~18,300 [20%] of those consultations are likely observable.) However, to maximize the utility of searches, users may focus effort on actions types with a combination of high observability and high consultation frequency, each of which can vary substantially (Figure 6). For example, formal consultations on road/highway projects has medium observability (0.48) but is a very common action type to be consulting on (*n* = 306 [with coordinates]). In contrast, new docks and piers in our sample had perfect observability, but there were only 34 such formal consultations.

**Figure 6.**
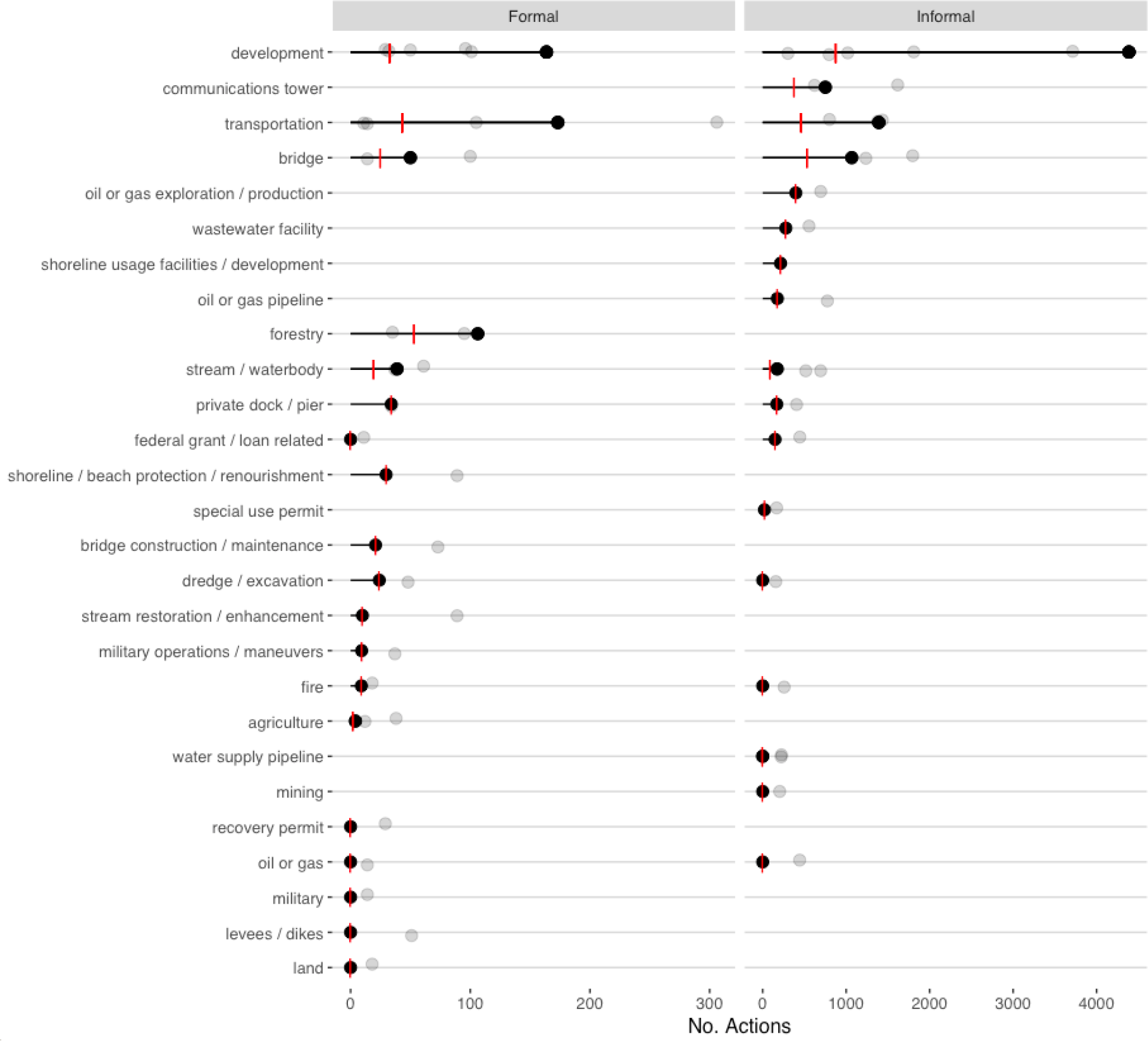
The combination of observability and number of consulted-on actions (“No. Actions”) help us find the most efficient categories of consultations to monitor. Lollipops show the total number of actions across work types, within work categories, we expect to find. (Notice separate scales for the formal and informal consultations.) Each gray circle is the number of actions for one or more work types within each work category. Red vertical bars are the mean expected number of actions per work type within each work category.

## Discussion

Compliance monitoring is an essential component of law and policy implementation. Even though section 7 is often considered the strongest regulatory component of the ESA, we do not currently know what the compliance “landscape” looks like. Are FWS assumptions of compliance—and therefore assumptions of the amount of harm to ESA-listed species they permit—realistic? Are the monitoring reports submitted by action agencies accurate? Freely available satellite and aerial images are one possible, cost-effective tool to help monitor compliance, but it was unknown how often these resources can be of use. Here we show that such resources may be a very effective component of section 7 compliance monitoring in some cases, but may be less useful in others.

We were able to observe in aerial images approximately 40% of 364 consultations from across 28 types of commonly consulted-on actions. Particular types of actions were more likely to be observed than others. For example, even though we assumed levee/dike work was likely to be observed, we could not find evidence for the three levee/dike actions we reviewed. In contrast, forestry actions had an observability of 0.82. In addition to variation between action types, there will likely be variation in observability between users. For example, our estimates of observability may be low relative to what FWS personnel would observe because of their greater familiarity with the projects. Even with variation—and under some simplifying assumptions, such as FWS recording geographic coordinates for all consultations—this average observability could translate to >4,000 FWS-approved actions per year for which it may be possible to monitor using freely available aerial or satellite imagery. This does not cover monitoring all consulted-on actions, but even partial coverage can be beneficial: holding some action agencies accountable may improve compliance in the broader community (Shimshack & Ward, 2005).

What are the barriers to using free imagery as a tool for section 7 compliance monitoring? First, FWS's lack of resources is a continuing challenge for the agency (GAO, 2003; Broderick, 2004; GAO, 2009; Gerber, 2016; GAO, 2017). They have insufficient resources to adequately conduct new consultations, much less spend time following up months or even years after the fact. In addition, there are likely institutional barriers to compliance monitoring by FWS. For example, reviewing monitoring reports is not prioritized in part because managers do not believe their responsibilities include “policing” action agencies that are also obliged to protect listed species (GAO 2009). Therefore, compliance monitoring will need to be a joint effort between governmental agencies and private citizens acting as watchdogs (Naysnerski & Tietenberg, 1992; Nie, 2008; GAO, 2017).

A second challenge is that compliance monitoring by people or groups other than FWS can only work if sufficient information is publicly available. In the current work we have focused on describing and quantifying our ability to find an action site given geographic coordinates, and to detect changes that are likely related to consulted-on actions. This is different from compliance monitoring, in which we need to know what FWS allowed or required under the terms of the consultations, then compare that to what was actually done. Unfortunately, FWS has not adopted transparency directives, such as Executive Order 13642 (Obama, 2013), to make their data publicly available. A handful of FWS offices post some formal consultation documents online, but gathering full documentation for most consultations would likely require voluminous Freedom of Information Act requests. Releasing full documentation for consultations may not be needed if FWS were to record authorized take as structured data, which has been recommended by the GAO (2006) and others (Totoiu et al., 2011; Clark, 2013; Evansen et al. 2017). But the agency has not taken steps to record take or release that data so that it can be used, e.g., to facilitate compliance monitoring. The integration of authorized take data and remotely sensed data is complicated because authorized take may be specified many different ways, such as the number of individuals taken or the amount of habitat that may be destroyed as “ecological surrogates” or “habitat proxies” (Totoiu et al. 2011; FWS 2015). In our experience, FWS biologists tend to specify take authorization in terms of habitat area to be affected more often than using individual measures. As such, current work practices at the agency may lend themselves to recording the data needed for remote sensing compliance monitoring.

### *Recommendations*

Based on our results and examining those results in the context of the “full stack” of needs for section 7 compliance monitoring, we make the following three recommendations:

1. *FWS needs to require that all consultations have accurate geographic coordinates recorded*. Because 48% of consultations recorded in TAILS lacked coordinates, there is near-zero chance of monitoring nearly half of consultations using remotely sensed imagery. FWS has very limited agency-wide data recording requirements, leaving decisions about which data should be recorded to individual regions or field offices (P. Ashfield,*pers. comm.)*. This shortcoming could be addressed immediately by requiring the coordinates field in TAILS be filled (and preferably bounds-checked) before FWS biologists submit a record.
2. *FWS needs to develop a system to record basic data about the size and scope of actions, including authorized take for formal consultations, and make those data publicly available*. While we were able to find many action sites, we did not perform actual compliance monitoring because we do not know what FWS had authorized. We do not know what FWS has authorized because they do not record that data systematically nor are those data released publicly. Calls to improve the tracking of authorized take have been advocated elsewhere (e.g., GAO, 2006; Totoiu, 2011; Clark, 2013; Evansen et al. 2017), and we echo those calls here. In addition to facilitating compliance monitoring— especially by conservation partners outside of FWS—the agency needs an authorized take database for their biologists to use in day-to-day work, e.g., to understand how much additional take may be authorized without harming chances of recovery. Beyond take of ESA-listed species (or their habitat) from formal consultations, recorded data needs to include a field for project footprint of all consultations.
3. *FWS should encourage their biologists to try using freely available satellite and aerial imagery for compliance monitoring as a normal part of section 7 implementation, and work with conservation partners to facilitate such monitoring*. We have shown that it is possible, using just geographic coordinates and very coarse action descriptions, to locate FWS-authorized actions using basic tools. Biologists from the agency have full access to the primary consultation documents and are familiar with their local areas; we expect they can be even more successful than we were at finding and following changes at action sites. But we suspect FWS biologists will be hesitant to try using recent technological advances such as freely available imagery without explicit direction, which senior staff can offer.

### *Conclusion*

Remote monitoring coupled with a more complete, open database can allow a multitude of average citizens to take part in protecting threatened and endangered species. For example, a surge in lawsuits against polluters in the 1980s may have been primarily due to accessible, relevant data that allowed students to easily flag Clean Water Act violations (Cohen, 1998). Private enforcement of the ESA currently revolves around citizen suit litigation against the FWS to challenge instances where the agency fails to carry out its responsibilities (Sobeck, 1993; GAO, 2017), but there may also be room for collaboration with FWS in enforcement actions against federal agencies or permittees. Satellite remote sensing will primarily be useful as a supplement to on-the-ground compliance monitoring (Purdy 2010). Recruiting online volunteers to flag suspicious action sites could allow FWS to efficiently allocate limited monitoring and enforcement resources in a targeted manner. The U.S. Environmental Protection Agency has adopted this very strategy through its Next Generation Compliance strategy. While most conservation civic science projects have involved monitoring of plants and wildlife (Theobald et al., 2015; Burgess et al., 2016; Sullivan et al., 2016), there are some precedents for using online applications to crowdsource the monitoring of illegal activities. For example, over 10,000 users on Tomnod (Baruch, May, & Yu, 2016) helped tag illegal forest fires in Indonesia (data here).

High-school students contributed to the detection of illegal deforestation in Borneo via the Earthwatchers project (See et al., 2016). As far as the authors know, the TAILS database used in this study could allow for the first opportunity to use an online crowdsourcing project to monitor compliance of endangered species regulations.

## Acknowledgements

We thank the hundreds of FWS biologists who diligently recorded data in TAILS for years; without their work this exploratory analysis would not have been possible. We also thank <NAMES> for useful feedback on earlier versions of this manuscript.

## Supplemental Materials

**SI Table 1. Expected observabilities for 448 action types recorded in the U.S. Fish and Wildlife Service consultation tracking database, TAILS.** Available at doi: 10.17605/OSF.IO/ANE85.

**SI Table 2. Observabilities across 364 consulted-on actions in 28 action types.** Available a1 doi: 10.17605/0SF.I0/ANE85.

**SI Figure 1.**
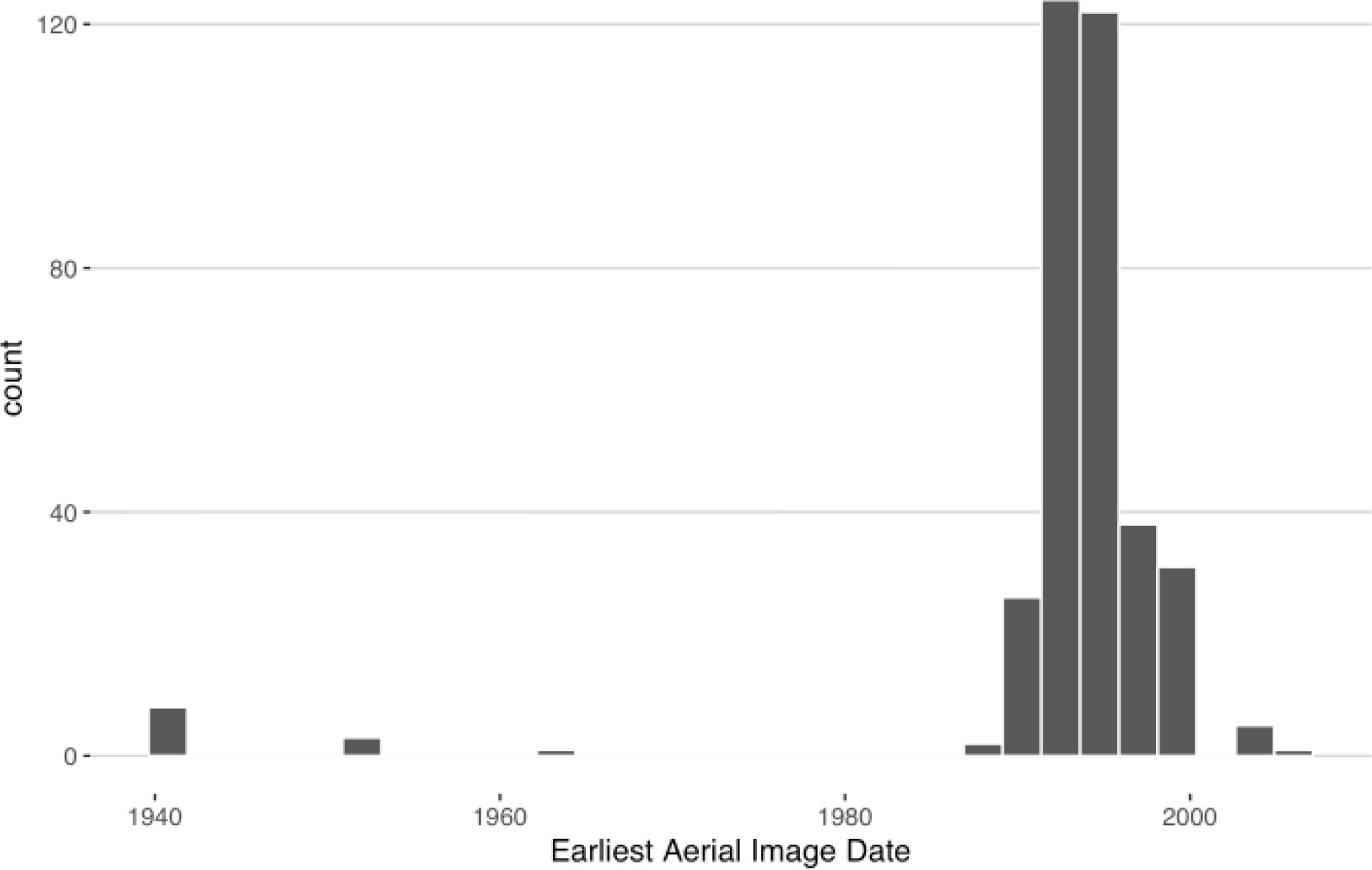
The distribution of the earliest available imagery dates from Google Earth Pro at the action sites we examined. Although some much older imagery was available in parts of Texas, most of the earliest images were from the 1990s.

